# PD-1 prelimits both the cytotoxic and exhaustion potential in thymic CD8+ T cells and impacts the maintenance of peripheral tumor immunity

**DOI:** 10.1101/2025.01.18.631253

**Authors:** Zhiming Mao, Jacob B. Hirdler, Joanina K. Gicobi, Mark Maynes, Michelle A. Hsu, Emilia R. Dellacecca, Wenjing Zhang, Jacob J. Teske, Ying Li, Geoffrey Zhao, Fabrice Lucien-Matteoni, Henrique Borges da Silva, Daniel D. Billadeau, Haidong Dong

## Abstract

Durable T cell immunity against cancer depends on the continual replenishment of effector CD8+ T cells. Thymic output has been correlated with favorable prognosis in cancer patients across a range of ages, suggesting that the thymus is an important source for replenishing T cells capable of controlling cancer progression. However, the effector potential of thymic mature CD8+ T cells and their regulation have not been clearly defined. In this study, we identified the ability of thymic single positive CD8+ T cells to gain effector potential after thymic selection, but they are subject to the regulation of PD-1. We found a previously undisclosed role of PD-1 in limiting both the cytotoxic and exhaustion potential of thymic and peripheral CD8+ T cells. Our results show that although PD-1 inhibition facilitates the expansion of effector CD8+ T cells, effector CD8+ cells gradually lose their antitumor activity within tumor tissues due to advanced exhaustion in the absence of PD-1. Thus, although the preset effector potential in thymic mature CD8+ T cells allows them to rapidly respond to malignant cells in the periphery, PD-1, as a checkpoint, is embedded in the thymic mature CD8+ T cells after positive selection to balance their effector function from exaggeration and exhaustion. Therefore, we propose that a strategy capable of upholding the cytotoxic capacity and avoiding exhaustion of CD8+ T cells during the early stages of PD-1 inhibition therapy is needed to achieve durable antitumor immunity.

## Introduction

Durable T cell immunity against cancer depends on the continual replenishment of cytotoxic CD8+ T cells that have been appropriately primed and differentiated for the elimination of cancer cells^1^. Recent studies have reported continued thymic output across a range of ages in humans and a strong correlation between thymic output and favorable prognosis in cancer patients, suggesting that the thymus is an important source for replenishing T cells capable of controlling cancer progression throughout a lifetime^2,3^. Before migrating to peripheral lymphoid organs, mature T cells must receive fate-determining TCR signals and undergo thymic positive and negative selection to ensure they are safe for the host and functionally competent^4,5^. Although the proliferation and migration competence of mature T cells in the thymus has been identified^5^, it remains unclear how the effector potential of CD8+ T cells is regulated in the thymus before they egress to peripheral tissues to mount durable control of cancer progression.

The success of immune checkpoint inhibitors (ICI) therapy targeting PD-1 or PD-L1 has been associated with the generation and expansion of cytotoxic CD8+ T cells T cells^6,7^. However, only a small portion of patients experience durable control of cancer progression after ICI therapy^8,9^. PD-1 inhibition is expected to revitalize effector CD8+ T cells from exhaustion, a functional state caused by persistent tumor antigen stimulation^10^. Current understanding is that PD-1 inhibition can restore or revitalize CD8+ T cells that are at stem-like or pre-exhaustion stage, but not those in terminal exhaustion stage^11,12^. However, it is still not clear whether PD-1 inhibition would affect the thymic output of mature CD8+ T cells capable of eliminating tumor cells in the periphery.

The role of PD-1 in thymic selection has been studied in the context of T cell receptor transgenic mice with germline PD-1 deficiency models^13^. Although the absence of PD-1 facilitates the beta-selection of thymocytes, the efficiency of positive selection is reduced without PD-1. This paradox is explained by the generation of double positive thymocytes due to enhanced beta-selection in the absence of PD-1, but these cells are not qualified for positive selection. This study seems to exclude a direct role of PD-1 in the process of positive selection but does not reveal how PD-1 affects the effector potential of positively selected thymic T cells due to the limitation of using global PD-1 deficient mice. Thus, it remains unclear whether PD-1 is involved in the regulation of the effector potential of thymic CD8+ T cells after positive selection in the thymus.

In this study, we used a CD8 (E8I)-specific conditional Pdcd1 knockout mice model to address the impact of PD-1 deficiency only at the stage after thymic positive selection. We found that PD-1 did not affect the outputs of positive selection but limited the positively selected single positive (SP) CD8+ cells from gaining cytotoxic potential by downregulating the expression of cytotoxic effector molecules (NKG7 and Granzyme B). Unexpectedly, we also found that PD-1 limited the exhaustion potential of SP CD8+ T cells by inhibiting the expression of TOX and TIM-3. In the periphery, the absence of PD-1 expanded the effector CD8+ T cells, which delayed tumor growth, but the advanced exhaustion potential of these effector CD8+ T cells eventually led to a loss of their antitumor activities. Our results imply that although PD-1 inhibition facilities the expansion of effector CD8+ T cells, the advanced exhaustion compromises the durable therapeutic effect of PD-1 inhibition. Therefore, a strategy capable of upholding the cytotoxic capacity and avoiding exhaustion of CD8+ T cells during early state of PD-1 inhibition therapy is needed to achieve durable antitumor immunity.

## Results

### PD-1 prelimits the cytotoxic and exhaustion potential of thymic single positive CD8+ T cells

The cytotoxic potential is one of the key functional features of CD8+ T cells. In the periphery, PD-1 regulates the cytotoxic potential of CD8+ T cells by suppressing actin remodeling at the immunological synapse and the release of cytotoxic granules^14^. In the thymus, the absence of PD-1 leads to enhanced beta-selection ^13^, but how PD-1 regulates the cytotoxic potential of single positive (SP) CD8+ T cells after thymic selection has not been clearly defined. To address this, we generated a mouse model with CD8+ T cell-specific Pdcd1 knockout (CD8 E8I-Cre-Pdcd1^fl/fl^, referred to as CD8-Pdcd1 cKO mice) (**Sup Fig 1A-B**), where the loss of Pdcd1 was induced in CD8+ T cells after their positive selection in the thymus, which was confirmed by flow cytometry (**Sup Fig 1F**). In this model, the absence of PD-1 did not affect the generation of SP CD8+ T cells based on their frequency compared to control mice (Pdcd1^fl/fl^, no CD8cre) (**Sup Fig 1C-E**).

**Figure 1.**
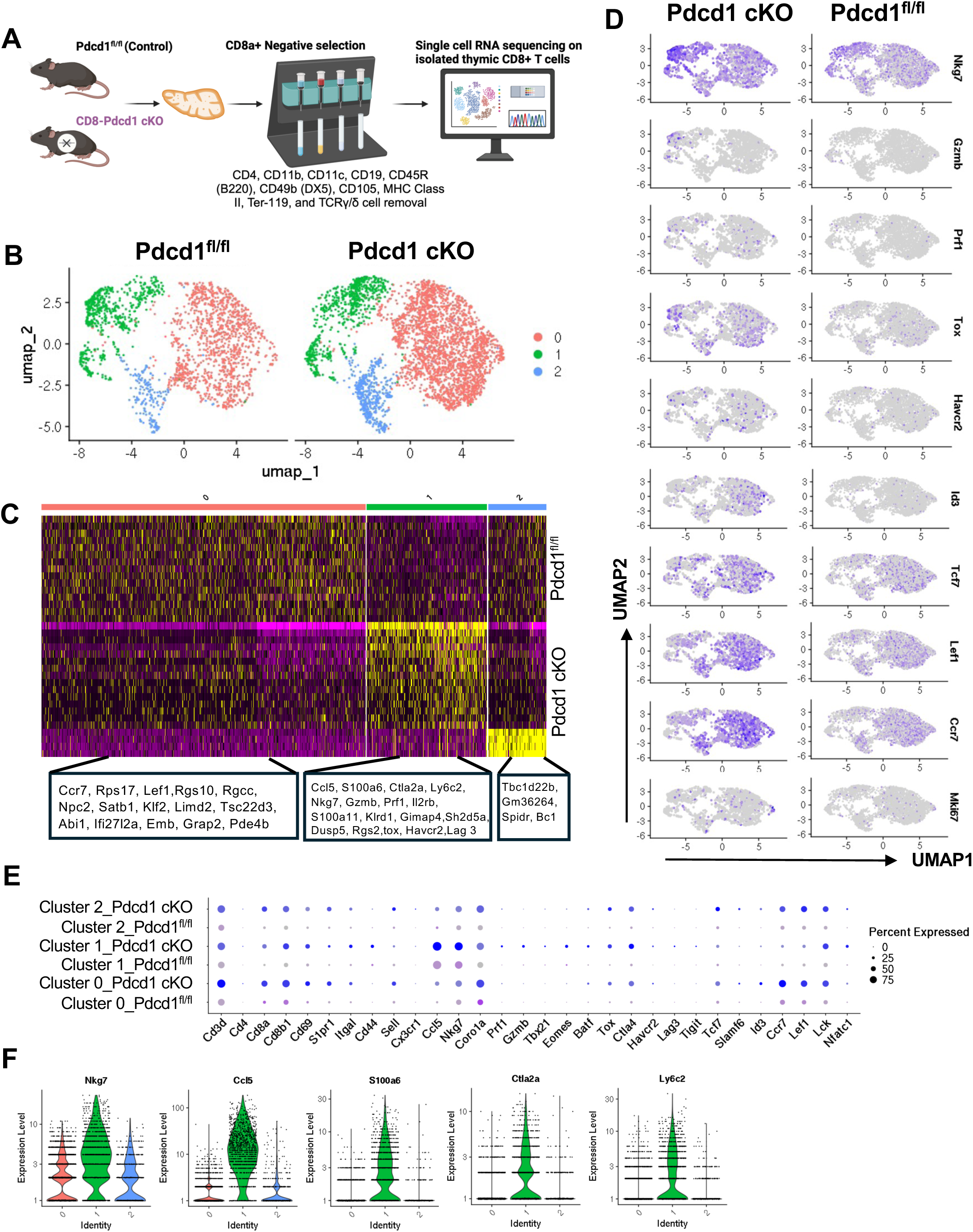
Increased transcription of genes encoding cytotoxic and exhaustion potential in thymic SP CD8+ T cells in the absence of PD-1. **(A)** Schematic of the isolation, processing, and single-cell RNA sequencing of thymic SP CD8+ T cells from Pdcd1^fl/fl^ (control) and Pdcd1^fl/fl^ CD8 E8I cre+ (CD8-Pdcd1 cKO) mice. **(B)** UMAP visualization of thymic SP CD8+ T cell clusters. **(C)** Heatmap of differential gene expression across three identified clusters. Yellow represents upregulated genes, while purple represents downregulated genes within each cluster. **(D)** Feature plots show the UMAP distribution and expression levels of *Nkg7*, *Gzmb*, *Prf1*, *Tox*, *Havcr2*, *Id3*, *Tcf7*, *Lef1*, *Ccr7*, and *Mki67* in thymic SP CD8+ T cells. Gene expression is depicted by a gradient, with darker blue representing higher expression levels. **(E)** Dotplot shows the expression of selected genes associated with cytotoxic effectors, exhaustion, stem-like features, and TCR signaling across the three clusters, stratified by control and CD8-Pdcd1 cKO mice. Dot color indicates the average expression level, with darker colors indicating higher expression (blue: CD8-Pdcd1 cKO; purple: control). Dot size represents the percentage of cells expressing each gene within a cluster. **(F)** Violin plots illustrate the expression levels of top candidate genes within each cluster across the three clusters. **(A)** is created using BioRender.

To examine whether PD-1 deficiency would affect the effector potential of SP CD8+ T cells in the thymus, we performed single-cell RNA sequencing (scRNA-seq) analysis of thymic mature SP CD8+ T cells isolated from control and CD8-Pdcd1 cKO mice (**Fig. 1A**). From there, we identified three cluster in UMAP graphs (**Fig. 1B**). Among them, we found Cluster 1 was enriched with genes encoding both cytotoxic and exhaustion molecules, which were more predominantly expressed in CD8-Pdcd1 cKO mice compared to control mice (**Fig. 1C-F**). This unique cluster 1 comprised genes encoding cytotoxic molecules (Nkg7, Prf1, Gzmb) and exhaustion markers (Tox, Ctla4, Havcr2 [Tim-3], Tigit, and Lag3), along with T cell activation-associated genes (Cd69, Lck, Nfatc1) (**Fig. 1E**). Other two clusters (cluster 0 and cluster 2) were identified with stem-like features, characterized by upregulated expression of genes such as Ccr7, Lef1, and Tcf7 (**Fig. 1E**). Notably, cluster 2 displayed additional expression of genes including Tbc1d22b, Gm36264, Spidr, and Bc1 (**Fig. 1C**). Our results suggest that PD-1 deficient SP CD8+ T cells gain an effector-like gene profile with both cytotoxic and exhaustion potential.

To validate our scRNA-seq analysis, we performed flow cytometry analysis of proteins associated with cytotoxic and exhaustion potential (**Fig. 2A**). We found that the frequency of NKG7+, GzmB+, and Perforin+ SP CD8+ T cells increased in the thymus of CD8-Pdcd1 cKO mice compared to control mice (**Fig. 2B-D**). Notably, two subsets of SP CD8+ T cells, recent positively selected semi-mature (TCRb+CD69+) and mature (TCRb+CD69-) cells, showed increased expression of NKG7+, Perforin+, and GzmB+ in the absence of PD-1 (**Fig. 2B-D**). Additionally, the transcription factors EOMES and T-Bet, which are associated with effector and memory differentiation, were also increased in SP CD8+ T cells in the absence of PD-1 (**Supp Fig. 1G-H**). In contrast, the stemness-associated transcription factor TCF-1 decreased in SP CD8+ T cells in the absence of PD-1 (**Fig. 2E**). Interestingly, we found that T cell exhaustion-associated molecules TOX and TIM-3 also increased in SP CD8+ T cells in the absence of PD-1 compared to control mice (**Fig. 2F-G**). Taken together, our results suggest that PD-1 limits thymic SP CD8+ T cells from gaining both the cytotoxic and exhaustion potential after thymic selection.

**Figure 2.**
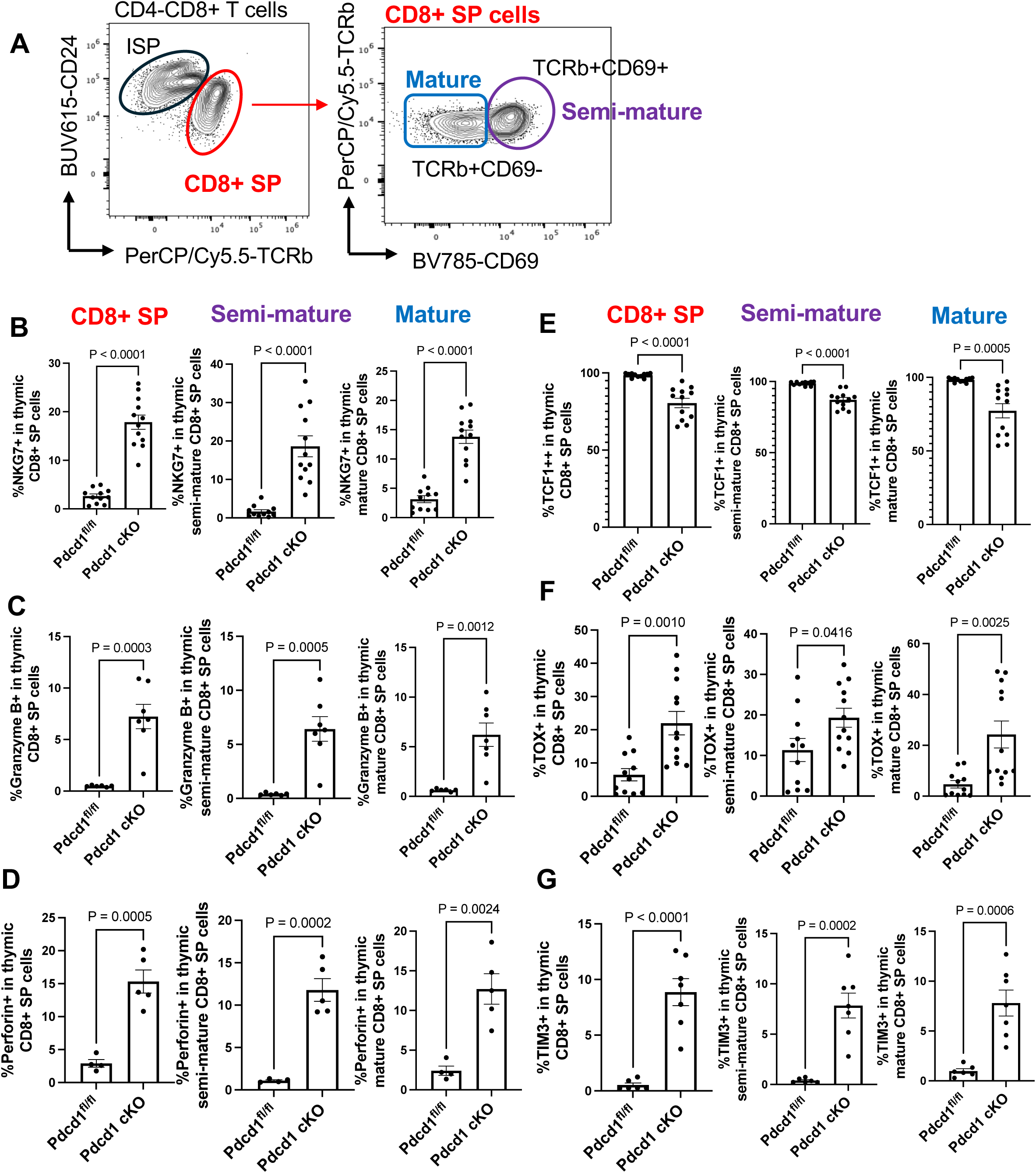
Increased protein expression of cytotoxic and exhaustion-associated molecules in thymic SP CD8+ T cells in the absence of PD-1. **(A)** Flow cytometry gating strategy for thymic SP CD8+ cells, identifying post-positively selected semi-mature (Tcrb+CD69+) and mature (Tcrb+CD69-) CD8+ T cell subsets. **(B-G).** Bar graphs show the frequency of NKG7 **(B)**, Granzyme B **(C)**, Perforin **(D)**, TCF-1 **(E)**, TOX **(F)**, TIM3 **(G)** in SP, semi-mature and mature thymic CD8+ T cells. P value was calculated using an unpaired, two-tailed t test. P values are shown at the top of each graph.

### Thymic single positive CD8+ T cells demonstrate cytotoxic effector function in the absence of PD-1

To determine the functional consequences of the increased expression of genes and proteins associated with cytotoxic function in SP CD8+ T cells in the absence of PD-1, we assessed degranulation (CD107a expression) and intracellular cytokine production (IFN-γ expression) following brief stimulation of isolated thymic SP CD8+ T cells with PMA and ionomycin. We found that degranulation increased in the effector-like TCF-1 SP CD8+ T cells in CD8-Pdcd1 cKO mice compared to control mice (**Fig. 3A**). Accordingly, the frequency of cytokine-producing cells also increased along with degranulation in CD8-Pdcd1 cKO mice compared to control mice (**Fig. 3C**). However, the expression of the exhaustion-associated molecule TOX does not seem to affect the cytotoxic capacity of SP CD8+ T cells in the absence of PD-1, as the frequency of TOX+CD107a+ cells increased in CD8-Pdcd1 cKO mice compared to control mice (**Fig. 3B**). To further understand the association of cytotoxic and exhaustion markers in SP CD8+ T cells, we measured the co-expression of TOX or TIM-3 with NKG7, a key molecule required for the cytotoxic capacity in CD8+ T cell killing ^15–17^. Unexpectedly, we found that NKG7 co-expressed with TOX or TIM-3 in thymic mature SP CD8+ T cells, and their frequencies increased in CD8-Pdcd1 cKO compared to control CD8+ T cells (**Fig. 3D-E**). Since NKG7 also co-expressed with GZMB in the absence of PD-1 (**Fig. 3F**), our results suggest that both the cytotoxic and exhaustion potential are coupled in the effector-like SP CD8+ T cells. However, the functional consequence of effector function seems to be decoupled from exhaustion potential in effector-like SP CD8+ T cells in the thymus.

**Figure 3.**
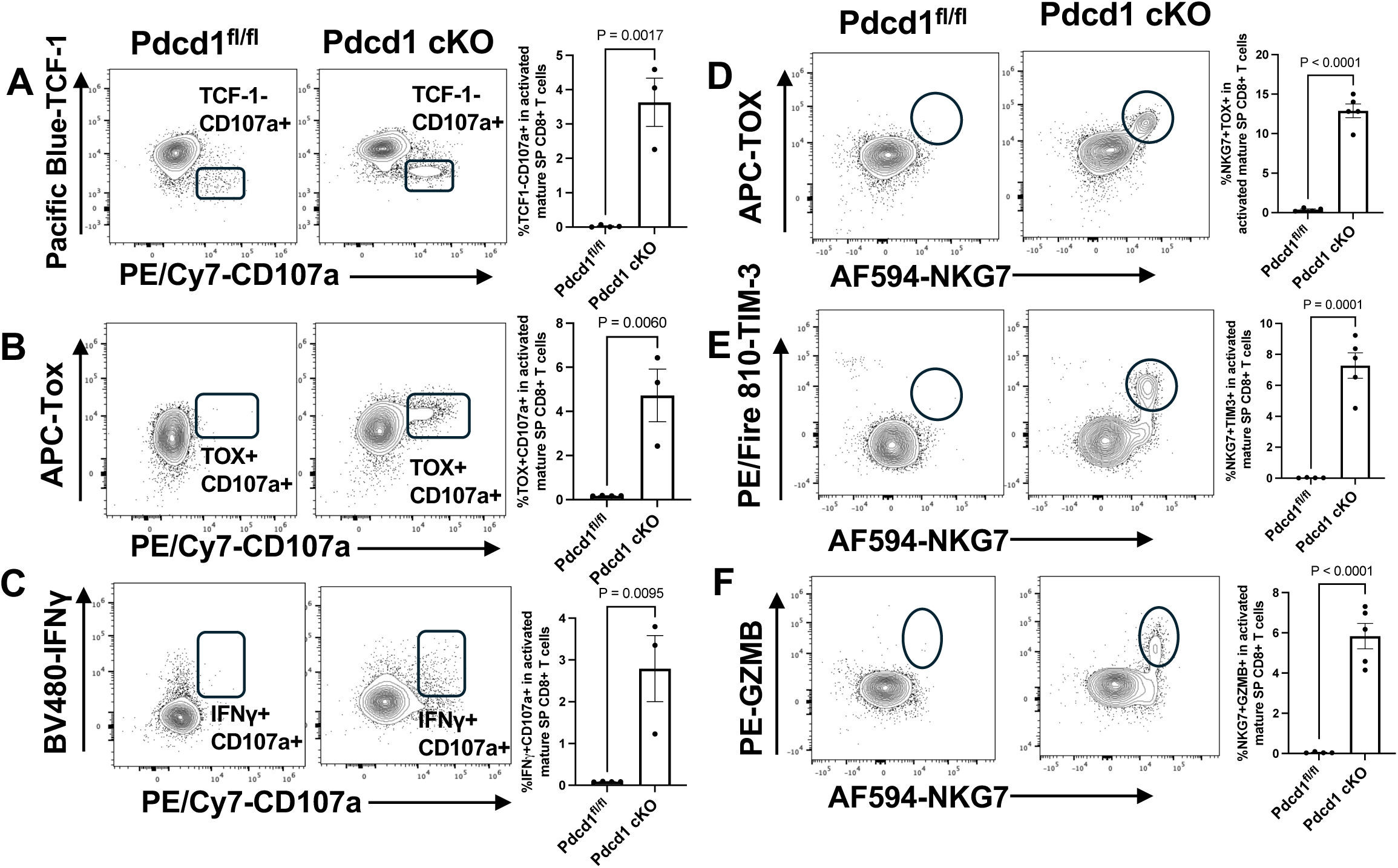
Increased functional cytotoxic capacities of thymic SP CD8⁺ T cells in the absence of PD-1. **(A-C)** Frequency of TCF-1- (**A**), TOX+ (**B**), IFNγ+ (**C**) along with CD107a+ in thymic SP CD8+ T cells after a 5-hour *ex vivo* stimulation with PMA and ionomycin. (**D-F**) Frequency of TOX+ (**D**), TIM-3+ (**E**), GzmB (**F**) along with NKG7+ in thymic SP CD8+ T cells. The representatives of gated population in dot plots are shown in the left and the summarized bar graphs are shown at the right. P value was calculated using an unpaired, two-tailed t test.

### Peripheral CD8+ T cells retained the cytotoxic and exhaustion potential post thymic selection in the absence of PD-1

To determine how the coupled cytotoxic effector and exhaustion potential in thymic CD8+ T cells would affect CD8+ T cells in the periphery, we measured and compared the phenotype and expression of molecules involved in cytotoxic and exhaustion potential in splenic CD8+ T cells. We found that CD8+ T cells with an effector phenotype (CD44+CD62L-) significantly increased in the spleen of CD8-Pdcd1 cKO mice compared to control mice (**Fig. 4A**). Accordingly, the effector CD8+ T cells expressed more cytotoxic molecules (NKG7, Perforin, and GzmB) as well as exhaustion-associated molecules (Tim-3 and TOX), but less TCF-1 in the absence of PD-1 (**Fig. 4B-G**). Additionally, the frequency of NKG7+GzmB+ cells increased in effector Pdcd1 cKO compared to PD-1^fl/fl^ CD8+ T cells (**Fig. 4H**). However, the expression of T-bet and the ratio of T-Bet/EOMES decreased within NKG7+GzmB+ cells in the absence of PD-1 (**Fig. 4I-J**). Our results suggest that although both the cytotoxic and exhaustion potential are preserved in peripheral effector CD8+ T cells in the absence of PD-1, the transcription factors of effector cells were downregulated in the peripheral effector CD8+ T cells.

**Figure 4.**
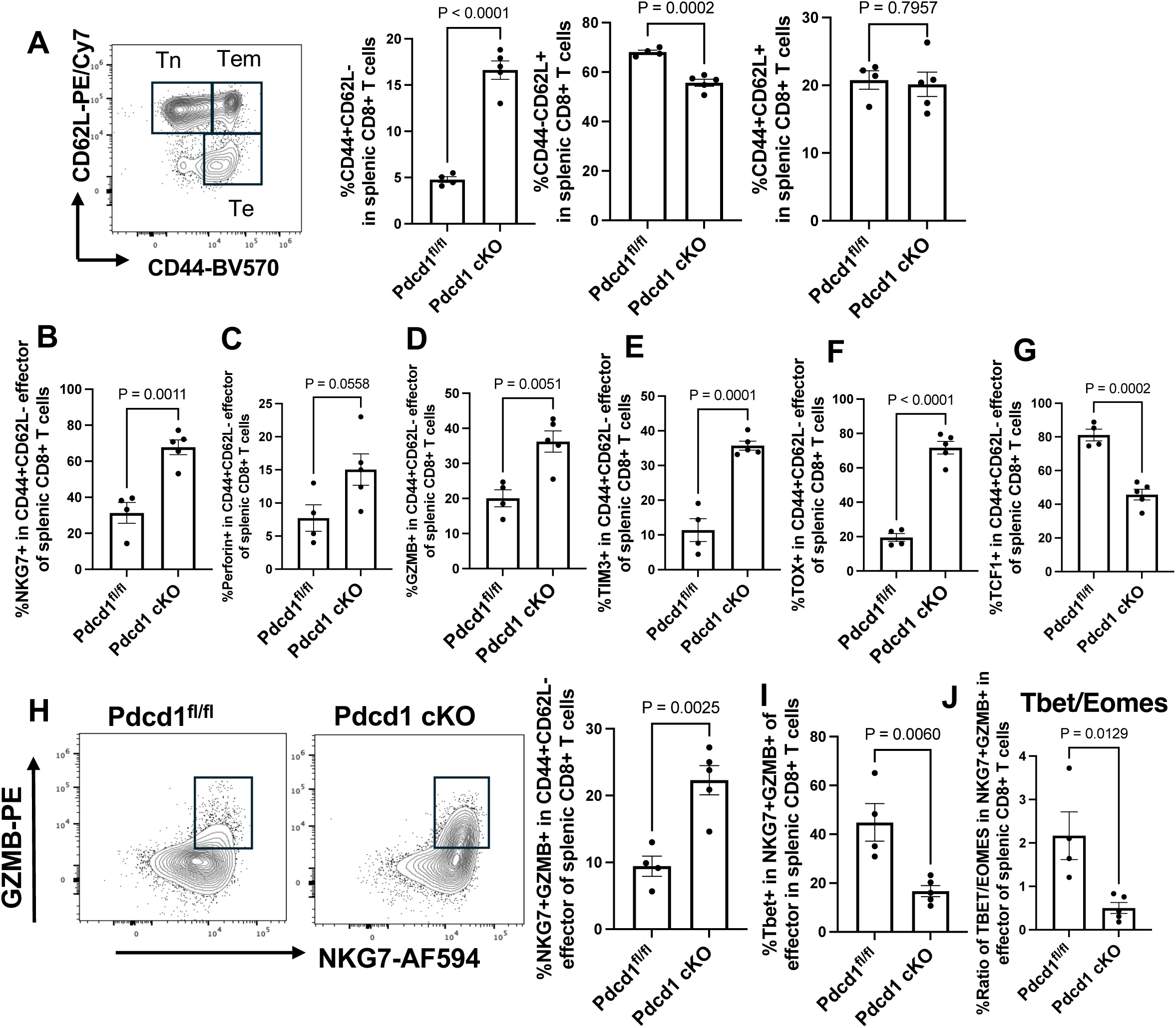
Increased cytotoxic and exhaustion potential of peripheral CD8+ T cells in the absence of PD-1. **(A)** Frequency of effector memory (CD44+CD62L+, Tem), effector (CD44+CD62L-, Te) and naïve (CD44-CD62L+, Tn) in gated TCRb+CD8+ T cells in the spleen. **(B-G)** The frequency of NKG7+ **(B)**, Perforin+ **(C)**, Granzyme B+**(D)**, Tim-3+ **(E)**, TOX+ **(F)**, TCF-1+ **(G)** in effector (CD44+CD62L-) splenic CD8+ T cells. **(H)** Frequency of NKG7+GZMB+ in effector (CD44+CD62L-) splenic CD8+ cells. **(I-J)** The percentage of T-bet+ (**I**) and the ratio of Tbet/Eomes (**J**) in NKG7+GZMB+ effector CD8+ T cells. P value was calculated using an unpaired, two-tailed t test.

In our scRNA-seq analysis of peripheral CD8+ T cells isolated from the spleens of CD8-PD-1 cKO mice and control mice, we identified six clusters in splenic CD8+ T cells (**Fig. 5A, Sup Fig 2A-C**). We noticed an increase in the percentage of cluster 5 in CD8-Pdcd1 cKO mice compared to Pdcd1^fl/fl^ (**Fig. 5B**). All clusters showed upregulated expression of Nkg7 in CD8-Pdcd1 cKO compared to control CD8+ T cells (**Fig. 5C-D**). However, only Cluster 5 was enriched with genes encoding both cytotoxic and exhaustion potential, such as Nkg7, Prf1, Gzmb, Havcr2 (Tim-3), Lag3, and Tigit (**Fig. 5C-D, Sup Fig 2B**). Notably, other cytotoxic molecules like Gzmb and Prf1, along with their transcription factors Tbx21 and Eomes, did not significantly appear in other clusters except Cluster 5 (**Fig. 5D, Sup Fig 2B**). Collectively, our results demonstrated that although peripheral CD8+ T cells carry over the effector phenotype and cytotoxic molecules at the protein level, the transcription of effector molecules (except Nkg7) is not active in peripheral CD8+ T cells in the absence of PD-1.

**Figure 5.**
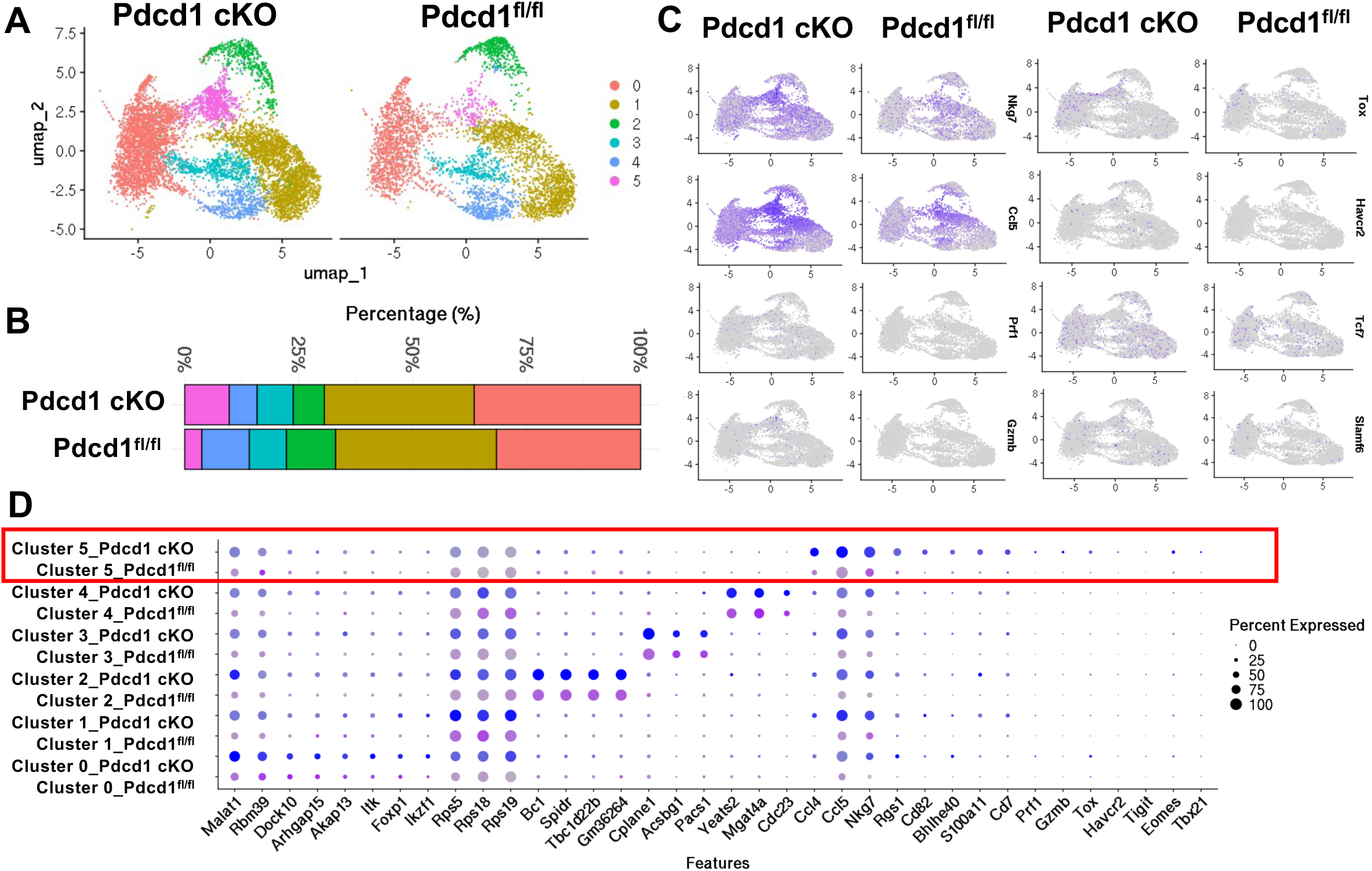
The immune profiles of splenic CD8+ T cells in the absence of PD-1. **(A)** UMAP visualization of clusters of CD8+ T cells in the spleen from control and CD8-Pdcd1 cKO mice. **(B)** Bar graphs show the percentage of each cluster within CD8+ T cells. (**C**) Feature plots show the UMAP distribution and expression levels of *Nkg7, Ccl5, Prf1*, *Gzmb*, *Tox*, *Havcr2*, *Tcf7*, and *Slamf6* in splenic CD8+ T cells. Gene expression is depicted by a gradient, with darker blue representing higher expression levels. **(D)** Dotplot graph shows the differential gene expression within each cluster and selected genes associated with cytotoxic capacity, exhaustion potential stratified by Pdcd-1 fl/fl and Pdcd-1 cKO CD8+ T cells.

### Tumor immunity is attributed to the expansion of cytotoxic CD8+ T cells but is lost due to the advanced exhaustion of these cells in the absence of PD-1

To determine whether the increased effector phenotype and cytotoxic molecules in peripheral CD8+ T cells would affect tumor immunity in the absence of PD-1, we measured and compared tumor growth between CD8-Pdcd1 cKO mice and control mice. We used three tumor models according to their degree of immunogenicity: MC38 (high), B16-OVA (intermediate), and B16F10 (low) (**Fig. 6A**). We found that most of the MC38 tumors (7/10) were rejected in CD8-Pdcd1 cKO mice compared to control mice (**Fig. 6B**). The growth of B16-OVA tumors was only delayed in CD8-Pdcd1 cKO mice, but with prolonged survival compared to control mice (**Fig. 6C**). However, the tumor growth of B16F10 tumors was not delayed or rejected and was comparable between CD8-Pdcd1 cKO mice and control mice (**Fig. 6D**). Our results suggest that the increased effector function of peripheral Pdcd1 cKO CD8+ T cells contributes to the control of tumors but is limited to tumors with high or intermediate immunogenicity.

**Figure 6.**
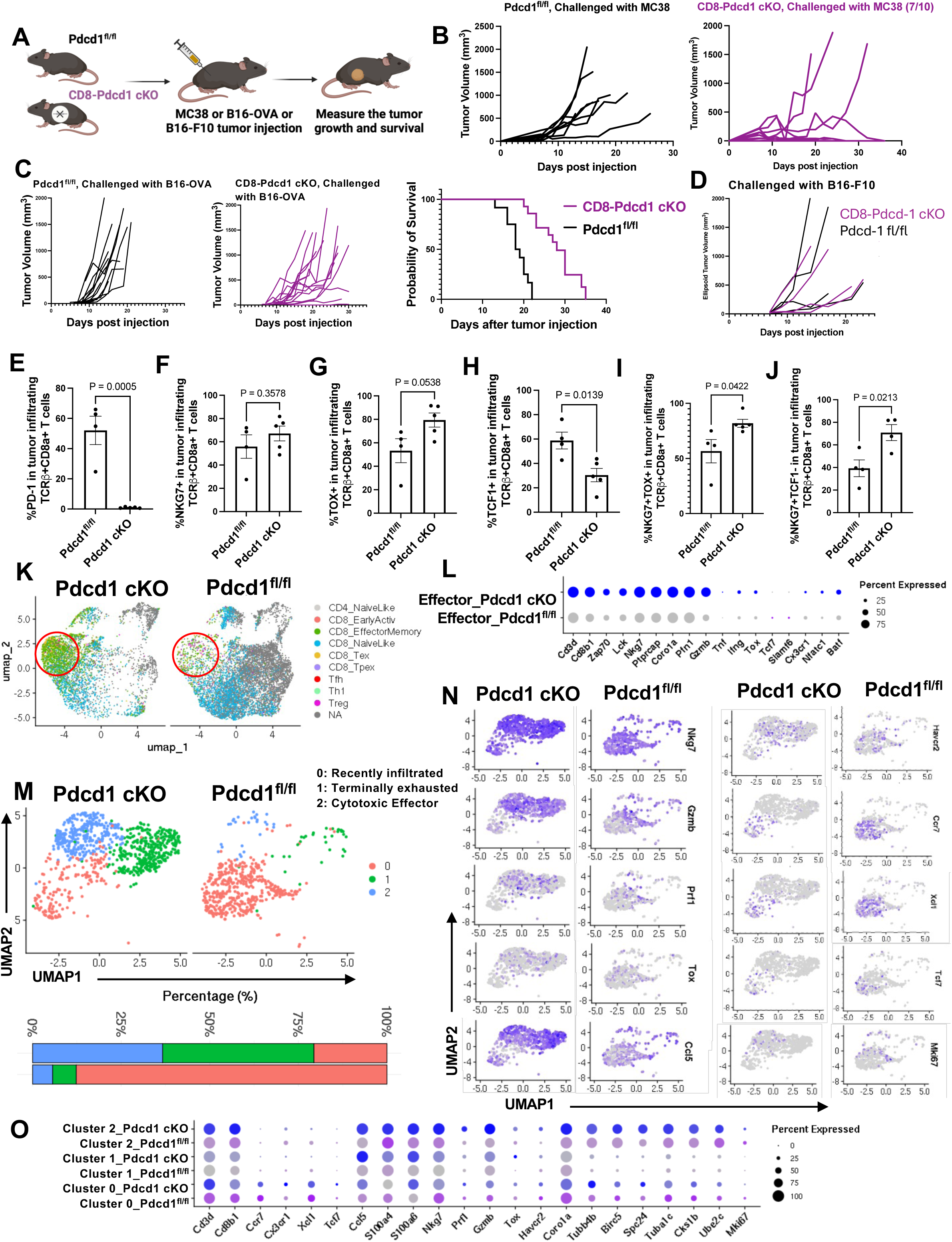
Tumor immunity is regulated by PD-1 according to the tumor immunogenicity. **(A)** Schematic of tumor models established in Pdcd1^fl/fl^ (control) and CD8-Pdcd1 cKO. **(B-D)** Tumor growth and survival curve of MC38 tumors (**B**), B16-OVA tumors (**C**), and B16-F10 tumors (**D**) in Pdcd1^fl/fl^ (control) and CD8-Pdcd1 cKO mice. (**E**-**J**) Frequency of PD-1+ **(E)**, NKG7+ **(F)**, TOX+(**G**), TCF-1+ (**H**), NKG7+TOX+ (**I**), NKG7+TCF-1-(**J**) in CD8+ TILs isolated at day 12 post B16-OVA tumor injection. **(K)** UMAP of mapped single-cell RNA-sequencing analysis of pooled CD8+ TILs (N=3) isolated on day 12 post B16-OVA tumor injection. **(L)** Dotplot graph shows the expression of genes encoding cytotoxicity and exhaustion in CD8+ TILs isolated as in (**E-J**). (**M**) Three main sub-clusters identified in the effector-like CD8+ TIL cluster from (**K**). (**N**) Feature plots show the UMAP distribution and expression levels of *Nkg7*, *Gzmb*, *Prf1*, *Tox*, *Ccl5*, *Havcr2*, *Ccr7*, *Xcl1*, *Tcf7* and *Mki67* in the sub-clusters of effector CD8+ TILs. Gene expression is depicted by a gradient, with darker blue representing higher expression levels. (**O**) Dotplot graph shows the signature genes within each sub-cluster for cluster identification stratified by control and CD8-Pdcd1 cKO mice. (**A**) is created using Biorender. P value was calculated using an unpaired, two-tailed t test.

To understand why the growth of tumors with intermediate immunogenicity was only delayed but could not be rejected in the absence of PD-1, we performed flow cytometry on the CD8+ TILs isolated from tumor-infiltrating lymphocytes on day 12 post B16-OVA tumor injection, a time point with peak accumulation of effector cells^18^. From there, we noticed a modest increase of NKG7+ CD8+ TIL in CD8-Pdcd1 cKO mice compared to control mice (**Fig. 6F**) in the absence of PD-1 (**Fig. 6E**), but the increase of TOX+, NKG7+TOX+, and NKG7+TCF-1-CD8+ T cells and the decrease of TCF-1 CD8+ T cells were significant in CD8-Pdcd1 cKO compared to Pdcd1^fl/fl^ CD8+ TILs (**Fig. 6G-J**). To further determine the immune profile of TILs in the absence of PD-1, we performed scRNA-seq analysis of CD8+ TILs isolated at day 12 post B16-OVA tumor injection. In comparison with our sequencing data and a previously published mouse TIL dataset^19^, we found that CD8+ TILs from CD8-Pdcd1 cKO mice exhibited a more pronounced “effector memory” phenotype compared to control mice (**Fig. 6K**). Among the CD8+ TILs, the genes encoding cytotoxic molecules Nkg7, Prf1, and Gzmb emerged as top candidate genes that were significantly upregulated in the effector/memory cluster of Pdcd1 cKO compared to Pdcd1^fl/fl^ CD8+ TILs (**Fig. 6L**). To understand how CD8+ TILs gradually lost their antitumor activity, we further sub-clustered the CD8+ TILs using our scRNA-seq data (**Fig. 6M, Sup Fig 3A**). From there, we identified two major clusters (Cluster 1 and Cluster 2) in Pdcd1 cKO CD8+ TILs and one major cluster (Cluster 0) in Pdcd1^fl/fl^ CD8+ TILs (**Fig. 6M**). Cluster 2 was characterized as “Cytotoxic effector CD8+ T cells” with elevated expression of genes encoding cytotoxic molecules, including Nkg7, Prf1, Gzmb, and Coro1a (**Fig. 6N-O**). Cluster 1 was characterized as “Terminally exhausted CD8+ T cells” with upregulated expression of Tox and downregulation of cytotoxic molecules including Nkg7, Prf1, Gzmb, and Coro1a (**Fig. 6N-O**). Of note, Cluster 2 also had a unique gene expression enriched with Tubb4b, Birc5, Spc24 compared to Cluster 1 (**Fig. 6O, Sup Fig 3B**). Cluster 0 was characterized as “Recently infiltrated CD8+ T cells” (Xcl1, Ccr7, Tafa2, Tcf7) (**Fig. N-O**). Therefore, our data suggest that the antitumor activity of CD8+ TILs may gradually decline due to advanced exhaustion in the absence of PD-1.

### NKG7 is subject to PD-1 regulation and is essential for PD-1 inhibition therapy

Our studies, as described above, show that the expression of NKG7 at both protein and transcription levels seems to be regulated by PD-1 in peripheral CD8+ T cells. Particularly, the suboptimal increase of NKG7 in CD8+ TILs may be associated with a loss of durable tumor immunity. To validate whether NKG7 expression is subject to PD-1 regulation in human tumor tissues, we performed spatial transcriptional analysis of NKG7+ vs. NKG7-CD8+ TILs in human cancer tissues **(Fig. 7A-D, Sup Fig 4A-B)**. Among four patients with muscle-invasive bladder cancer (**Fig. 7A, Sup Fig 3A**), we were able to select regions of tissue enriched with NKG7+ CD8+ T cells or NKG7-CD8+ T cells (**Fig. 7B**). From there, we found a different list of genes that were upregulated in NKG7+ CD8+ T cells vs. NKG7-CD8+ T cells (**Fig. 7C**). Pathway analysis indicated that NKG7-CD8+ T cells, but not NKG7+ CD8+ T cells, were enriched with genes positively associated with the “PD-1/PD-L1 cancer immunotherapy pathway” (**Fig. 7D**), suggesting that the PD-1/PD-L1 pathway may limit the expression of NKG7 in NKG7-CD8+ TILs in human tumor tissues.

**Figure 7.**
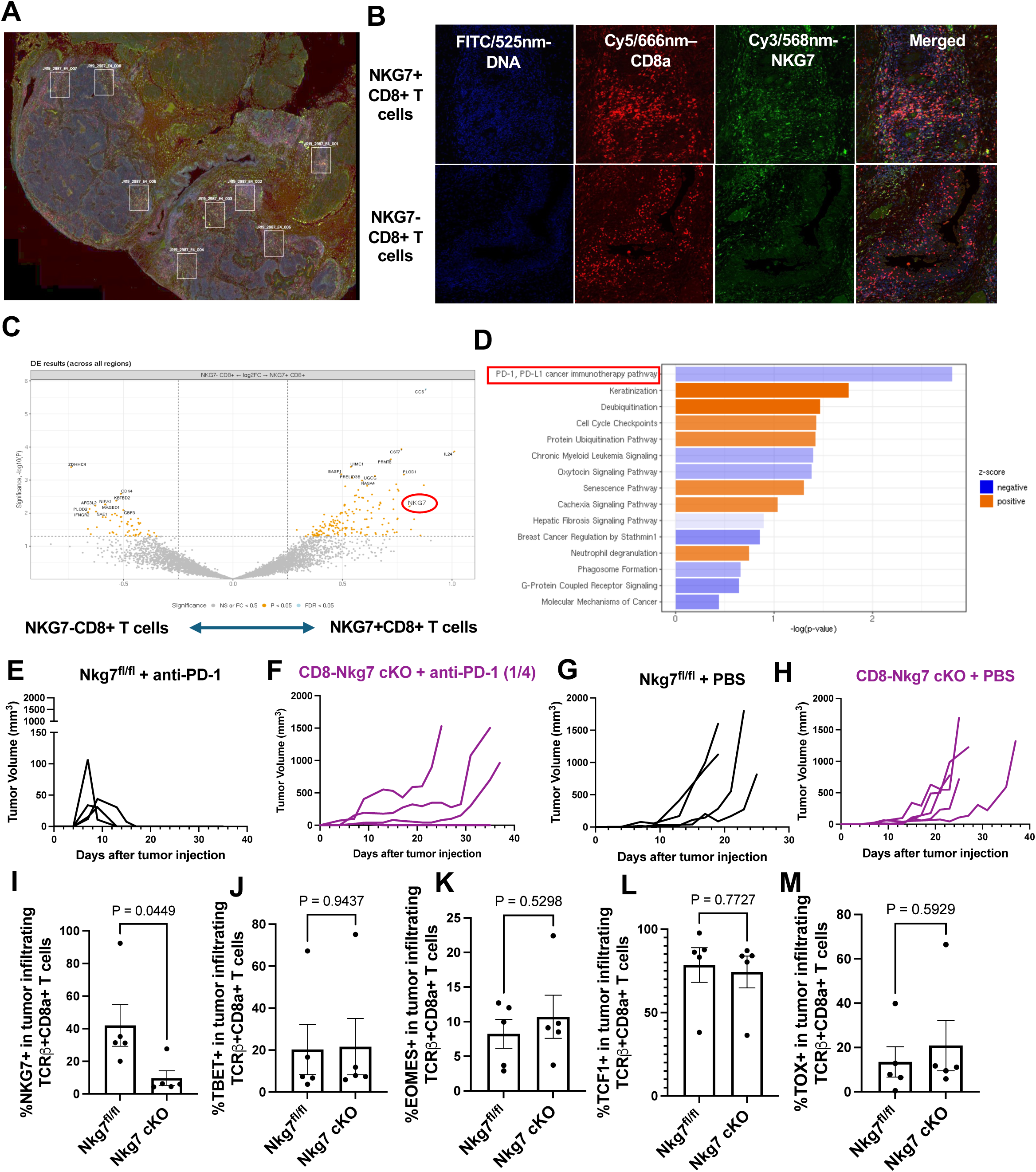
NKG7 is subject to PD-1 regulation and essential for PD-1 inhibition therapy. **(A)** Representative regions of Interests (ROI) for spatial transcriptome analysis were selected from tissue slides of patients with muscle invasive bladder cancer. (**B**) Representative images of NKG7+CD8+ T cells (Up) and NKG7-CD8+ T cells (Below) with corresponding staining. **(C)** Differential Gene Expression (DEG) analysis of the changes of gene expression in NKG7+CD8+ T cells vs. NKG7-CD8+ T cells. NKG7 was highlighted in the plot. (**D**) Pathway enrichment analysis shows the top pathway (PD-1, PD-L1 pathway, boxed) in association with NKG7-CD8+ T cells (Z score negative, Blue), but not with NKG7+CD8+ T cells (Z score positive, Orange). (**E**-**H**) Tumor growth of MC38 tumors treated with anti-PD-1 (**E-F**) or PBS (**G-H**) in Nkg7^fl/fl^ (Control) and CD8-Nkg7 cKO mice. (**I**-**M**) Frequency of NKG7+ (**I**), T-bet+ (**J**), EOMES+ (**K**), TCF-1+ (**L**), TOX+ (**M**) in CD8+ TILs isolated at day 14 post MC38 tumor cell injection. P value was calculated using an unpaired, two-tailed t test.

To determine whether NKG7 is necessary for CD8+ TILs to eliminate tumors in response to PD-1 inhibition, we generated CD8-specific Nkg7 cKO mice in which MC38 tumors were treated with anti-PD-1 antibody. As expected, anti-PD-1 therapy completely rejected the MC38 tumors in control (Nkg7^fl/fl^) mice but only partially rejected tumors in CD8-Nkg7 cKO mice (**Fig. 7E-F**). Since the tumor growth was comparable between control and CD8-Nkg7 KO mice without anti-PD-1 treatment (**Fig. 7G-H**), the loss of NKG7 does not seem to cause immunodeficiency, which was confirmed by the comparable expression of transcription factors involved in stem-like (TCF-1), effector (T-bet/Eomes), or exhaustion (TOX) in Nkg7 cKO vs. Nkg7^fl/fl^ CD8+ TILs (**Fig. 7J-M**) in the absence of NKG7 (**Fig. 7I**). Thus, our results indicate that NKG7 is required for PD-1 inhibition to motivate effector CD8+ TILs to eliminate tumors.

### The expansion of effector CD8+ T cells is transient in the absence of PD-1, but a sequential blockade of PD-L1 can sustain antitumor immunity

The less regression of B16-OVA tumors compared to MC38 tumors suggests that highly immunogenic tumors may rapidly expand effector cells in the absence of PD-1 before the exhaustion program takes over the functional state of CD8+ T cells in tumor tissues. To test whether a rapid expansion of effectors in the absence of PD-1 would reject less immunogenic tumors (B16-OVA), we immunized CD8-Pdcd1 cKO mice and control mice with OVA protein and poly(I:C) (as an adjuvant) and challenged the immunized mice with B16-OVA tumor cells on day 7 after immunization (**Fig. 8A**), a time with peak accumulation of effector cells. As expected, we found a complete rejection of B16-OVA tumors in CD8-Pdcd1 cKO mice compared to control mice (**Fig. 8B**). Interestingly, we found that a long-term tumor-specific memory was established in the CD8-Pdcd1 cKO mice after they completely rejected the tumor on day 7 with rapidly expanded effector cells, as they rejected the second challenge of B16-OVA tumors but not B16F10 tumors (**Fig. 8C**). However, when the effector cells gradually declined over time on day 21 after immunization, only a partial rejection of B16-OVA tumors was found in CD8-Pdcd1 cKO mice (**Fig. 8D**). Accordingly, our flow cytometry analysis showed a rapid accumulation of effector cells (identified by CD44+CD62L-, CX3CR1+ and Cd107a+) peaking on day 7 (week 1) after immunization in CD8-Pdcd1 cKO mice compared to control mice but followed with a decline of these effector cells thereafter (**Fig. 8E-G**).

**Figure 8.**
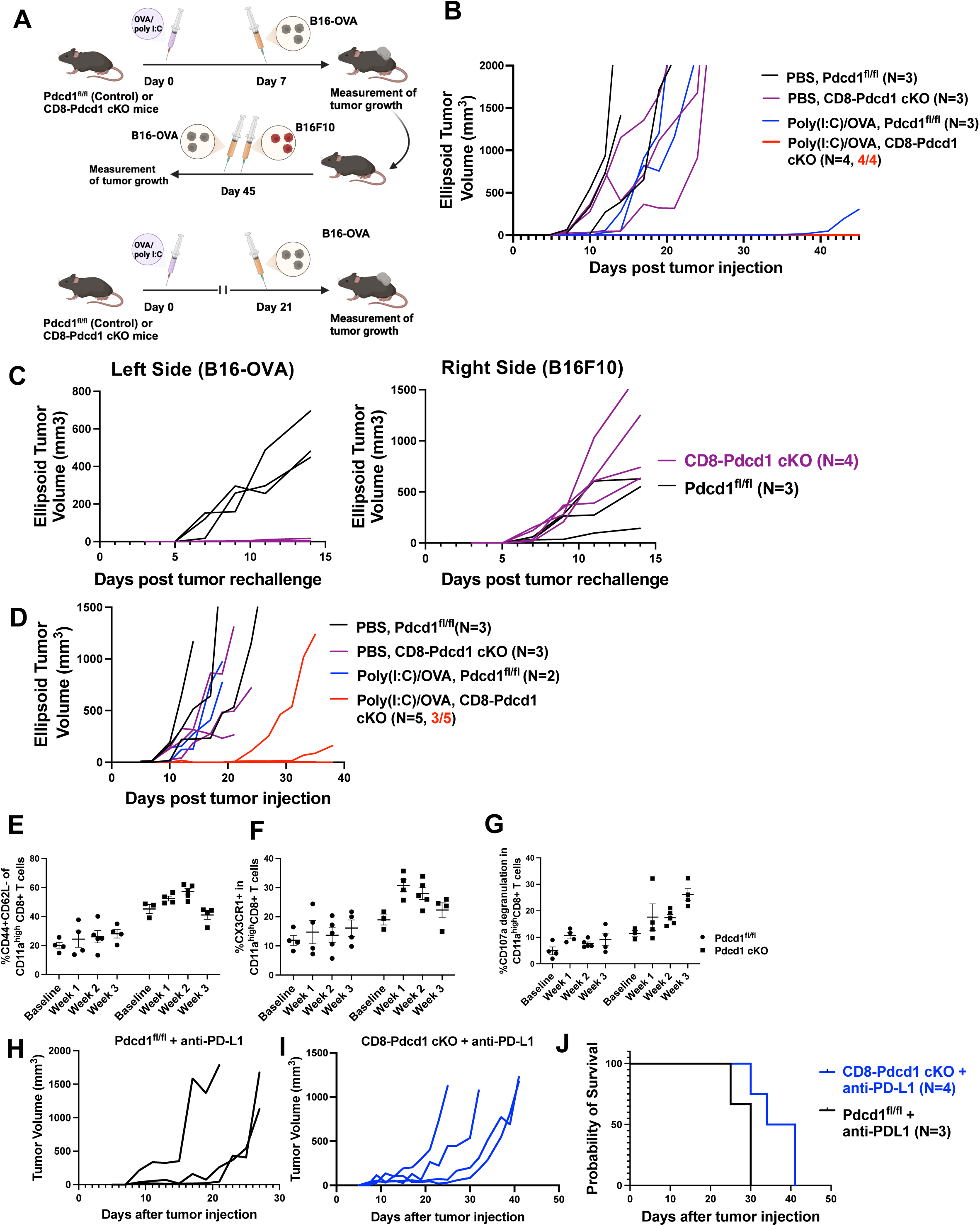
Transient expansion of effector CD8+ T cells in the absence of PD-1. **(A)** Schematic of tumor challenges in mice immunized with OVA/ poly(I:C). **(B)** The growth of B16-OVA tumors after tumor injection on day 7 after immunization. (**C**) The growth of B16-OVA (left flank) or B16F10 (top right flank) tumors injected on day 45 in mice that rejected the first tumor challenge at lower right flank on day 7 as in (**B**). The growth of B16-OVA or B16F10 tumors in naïve mice was used as control. **(D)** The growth of B16-OVA tumors after tumor injection on day 21 after immunization. (**E**-**G**) Frequency of CD44+CD62L+(**E**), CX3CR1+(**F**), CD107a degranulation (**G**) in splenic CD11a^high^ CD8+ T cells from the baseline and weeks after immunization. CD11a was used to identify antigen-primed T cells. (**H-J**) the tumor growth (**H-I**) and survival (**J**) of mice with B16-OVA tumors after treatment with anti-PD-L1 (10B5) starting at day 7 after tumor injection for a total of five doses.

Next, we tested whether a sequential blockade of PD-L1 would further expand effector cells in the absence of PD-1. Besides binding to PD-1, PD-L1 binds CD80, which is required to stimulate CD28 for costimulatory signals needed for CD8+ T cells to avoid exhaustion^20^. We produced an anti-PD-L1 antibody (10B5), which can block the interaction of PD-L1 with PD-1 and CD80^21^. We expected the 10B5 antibody would allow more CD80 to stimulate CD28 by releasing CD80 from binding with PD-L1, thereby expanding effector cells through co-stimulation and avoiding exhaustion. To that end, we found that 10B5 antibody treatment further delayed the tumor growth of less immunogenic B16-OVA tumors in CD8-Pdcd1 cKO mice compared to control mice, along with prolonged survival of the mice (**Fig. 8J-L**). Taken together, our results indicate that while the absence of PD-1 facilitates the expansion of cytotoxic CD8+ T cells, PD-1 deficient CD8+ TILs gradually lose their antitumor activity within tumor tissues due to the advanced exhaustion (**Sup Fig. 5**). Therefore, a strategy capable of upholding the cytotoxic capacity and avoiding exhaustion of CD8+ TILs is needed to achieve durable antitumor immunity.

## Discussion

Our results show that the ability of CD8+ T cells to gain both cytotoxic and exhaustion potential happens earlier in the thymus, but they are subject to the regulation of PD-1. Our findings suggest that the effector potential could be a preset feature of SP CD8+ T cells after thymic positive selection, before they encounter cognate antigens in the periphery. This preset effector potential would allow CD8+ T cells to rapidly acquire effector function in the periphery when they encounter pathogens or malignant antigens, thereby protecting the host from infections and tumors. However, this also raises the risk of normal tissue damage if their effector function is exaggerated. To this end, PD-1, as a checkpoint, is embedded in the selected SP CD8+ T cells to prevent them from exaggerating their function.

An unexpected finding from our study is the role of PD-1 in limiting the exhaustion potential in SP CD8+ T cells, challenging the dogmatic view of PD-1’s role in T cell exhaustion in the periphery. While the primary role of PD-1 is to restrain CD8+ T cells from gaining effector function, the loss of PD-1 also promotes effector cells to advance into exhaustion. Our observation aligns with a previous report showing that PD-1 signals prevent effector T cells from becoming exhausted in peripheral tissues^22^. Thus, the dual role of PD-1 provides an elegant balance of the magnitude of effector CD8+ T cells in response to antigen stimulation. To this end, we found that although the absence of PD-1 allows a rapid expansion of effector CD8+ T cells to reject highly immunogenic tumors, the advanced exhaustion in effector CD8+ T cells in the absence of PD-1 compromises the control of tumors with less or no immunogenicity.

The field of anti-PD-1 therapy is evolving from monotherapy to combination therapy designed to further expand effector T cells. However, it is not completely clear which pathway would allow us to design an optimal combination strategy that can synergistically work with anti-PD-1 blockade to establish durable antitumor immunity. To this end, distinguishing the role of PD-1 in the regulation of cytotoxic and exhaustion potential in CD8+ T cells would provide a pathway for designing rational combination therapy. Based on our discovery of PD-1’s role in limiting both the cytotoxic and exhaustion potential of CD8+ T cells in thymic and peripheral tissues, a strategy that promotes effector function while preventing exhaustion should be implemented early in anti-PD-1 therapy.

Leveraging an anti-PD-L1 antibody that can block both PD-1 and CD80 ^21^, we found that this anti-PD-L1 antibody can further delay tumor growth in the absence of PD-1. The potential underlying mechanism could be that, in the absence of PD-1, the anti-PD-L1 antibody mainly blocks the binding of PD-L1 with CD80, allowing more CD80 to stimulate CD28. This provides costimulatory signals via CD28 for effector CD8+ T cells to expand and avoid exhaustion^20^. From there, more preclinical and clinical studies are expected to develop strategies aimed at integrating both PD-1 inhibition and CD28 co-stimulation to achieve durable tumor immunity.

Although the use of the E8I-cre system to identify the cytotoxic potential in single positive (SP) CD8+ T cells represents a critical first step in the mechanistic analysis of how the effector capacity of CD8+ T cells is preset during T cell development, additional work is needed. For example, it is not clear how PD-1 signals are integrated into the regulation of both the cytotoxic and exhaustion programs in SP CD8+ T cells during or after thymic selection. Additionally, it remains to be determined whether TCR signals are the initial signals in presetting the cytotoxic or exhaustion potential in thymic CD8+ T cells.

From there, we must investigate whether the transcription of genes involved in cytotoxic and exhaustion potential can be decoupled in thymic or peripheral CD8+ T cells through epigenetic regulation. Since PD-1 signals mainly regulate the downstream pathway of CD28, we propose that enhancing CD28 co-stimulatory signals may be an option to tip the balance of cytotoxic and exhaustion potential in CD8+ T cells. In this direction, we found that using a PD-L1 antibody that can block the binding of PD-L1 and CD80, allowing more CD80 to engage CD28, could be a promising strategy to enhance CD28 signals.

In summary, our study identified a previously undisclosed role of PD-1 in limiting both the cytotoxic and exhaustion potential of thymic and peripheral CD8+ T cells. Our results show that although PD-1 inhibition can expand effector CD8+ T cells, advanced exhaustion may compromise the durable antitumor activity of these cells. Thus, mere PD-1 inhibition may only provide transient control for tumors with weak or low immunogenicity. Consequently, our study provides a direction for designing a synergistic strategy that can expand effector CD8+ T cells and prevent exhaustion at the early stages of PD-1 inhibition therapy.

## Material and Methods

### Mouse

Pdcd1^flox/flox^ ^(fl/fl)^ mice were a kind gift from Dr. Vassiliki A. Boussiotis from Harvard Medical School, and the protocol for generating these mice has been previously described^23^. Nkg7^fl/fl^ mice were generated by inGenious Targeting Laboratory (Ronkonkoma, NY). E8I CD8-Cre mice (Strain #: 008766) were purchased from the Jackson Laboratory. The Pdcd1^fl/fl^ E8I CD8Cre (CD8-Pdcd1 cKO) mice were generated by crossing Pdcd1^fl/fl^ mice with E8I CD8-Cre mice. The Nkg7^fl/fl^ E8I CD8Cre (CD8-Nkg7 cKO) mice were generated by crossing Nkg7^fl/fl^ mice with E8I CD8-Cre mice. Pdcd1^fl/fl^ mice were genotyped using the primer pair: 1 – 5’ TAT CCC TGT ATT GCT GCT GCT G 3’ and 2 – 5’AAT GAA TTG AGG AGT AGG GCC TG3’. E8I CD8Cre were genotyped using the following primer 1: 5’ CAA TGG AAG GAA GTC GTG GT 3’. Primer 2: 5’ TGG GAT TTA CAG GGC ATA CTG 3’ and primer 3: 5’ CAC ACA TGC AAG TCT AAA TCA GG 3’. Pdcd1^fl/fl^ mice were genotyped using primer pair 1: 5’-GTG GCA CAG GAT CTC TCA GG -3’. 2: 5’- GAT CAA GGG CTT GCC TTA GC- 3’. Animals were maintained under specific-pathogen-free conditions at Mayo Clinic Rochester. In all experiments, male and female mice used were after 8-12-week-old and were randomly assigned for experiments. All experimental procedures were approved by the institutional animal care and use committee at Mayo Clinic Rochester (IACUC: A00006353-21-R24).

### Tumor Cells, Media, and Cell Culture

Dulbecco’s modified Eagle medium (DMEM, Gibco, Cat#11885-084), DMEM with 4.5 g/L glucose, L-glutamine and sodium pyruvate (DMEM^high^, Corning, Cat #10-013-CV) and RPMI 1640 (Corning, Cat#10-040-CV) were supplemented with 10% heat-activated fetal bovine serum (FBS, Gibco, Cat # A52567-01) and 10 mM HEPES buffer (Corning, Cat # 25-060-CI) and 1X Penicillin/Streptomycin (Cellgro, Cat#30-002-CI) for a complete medium. MC38 colon adenocarcinoma cells were purchased from MIlliporeSigma (SCC172) and has been maintained using DMEM^high^ complete medium. B16-F10 murine melanoma cell line was purchased from ATCC (CRL-6475) and cultured using DMEM complete medium. B16-OVA murine melanoma cell line was a gift from Dr. Richard Vile at Mayo Clinic Rochester and cultured using RPMI 1640 complete medium with 0.8 mg/mL Geneticin™ Selective Antibiotic (G418 Sulfate, Gibco, Cat#11811031). Cell lines were screened for *Mycoplasma* contamination and authenticated by short tandem repeat profiling (B16 at American Type Culture Collection and MC38 at IDEXX).

### Tumor models and *in vivo* treatment

Pdcd1^fl/fl^ E8I CD8Cre- (control) and Pdcd1^fl/fl^ E8I CD8Cre+ (CD8-Pdcd1 cKO) mice or Nkg7^fl/fl^ E8I CD8Cre- (control) and Nkg7^fl/fl^ E8I CD8Cre+ (CD8-Nkg7 cKO) were subcutaneously injected with 0.5 million tumor cells resuspended in 100μL PBS. Tumor growth was measured every 2-3 days starting from day 7 post tumor injection until the euthanasia endpoint in compliance with animal care guidelines. Perpendicular tumor diameters were measured using a digital caliper (Carbon Fiber Composite, Fisher Scientific). Tumor size was calculated as ellipsoid tumor volume using the formula: 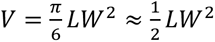 . For immune checkpoint blockade studies, control and CD8-Pdcd1 cKO mice were intraperitoneally injected with 200 μg anti-PD-L1 antibody (Clone 10B5, Sigma-Aldrich, Cat# MABC1131-100UG) starting from day 7 post tumor injections every two days for a total of 5 doses. Control and CD8-Nkg7 cKO mice were intraperitoneally injected with 200μg anti-PD-1 antibody (Clone G4, Sigma-Aldrich, Cat # MABC1132-100UG). For Ovalbumin (OVA) immunization studies, 50μg polyinosinic-polycytidylic acid (Poly(I:C), Novus, Cat#NBP2-25288) and 500μg OVA peptide (MilliporeSigma, Cat# A5503) were mixed in 200 μL PBS and intraperitoneally injected to the mice.

### Isolation of mouse CD8+ T cells

Mouse splenic and thymic CD8+ T cells were isolated using CD8a+ T cell isolation kit (Miltenyi Biotec, Cat#130-104-075) according to manufacturer instructions. For isolating CD8+ tumor infiltrating lymphocytes (TILs), tumors were first cut into small pieces of 2-4mm and digested using tumor dissociation kit at 37°C for 45 minutes with gentle agitation (Miltenyi Biotec, Cat#130-096-730) according to manufacturer instructions. Dead cells were further removed using Dead cell Removal kit (Miltenyi Biotec, Cat# 130-090-101). CD8+ TILs were then isolated using CD8 (TIL) microbeads, mouse (Miltenyi Biotec, Cat#130-116-478) based on manufacturer instructions.

### Single cell RNA sequencing and analysis

Mouse thymic CD8+ T cells, splenic CD8+ T cells and CD8+ TILs were isolated from adult mice (>8 weeks old) for single-cell RNA sequencing. Single cell libraries were prepared using Fluent Biosciences Pre-Templated Instant Partitions Sequencing (PIPseq) v3 Kit according to the manufacturer’s protocol. Briefly, cells were first filtered through a 40-μm cell strainer for the debris removal. The viability was assessed through Trypan Blue cell counting, and samples with more than 80% viability were proceeded for sample preparation. Each sample was loaded to the PIPseq microfluidic system, which partitioned individual cells into droplets containing molecular barcodes for unique transcript identification. During reverse transcription, unique molecular identifiers (UMIs) and well-specific barcodes were incorporated to enable precise quantification and tracking of individual transcripts. Subsequently, cDNA synthesis and amplification were performed to maximize transcript coverage and minimize bias. Amplified cDNA was purified and quantified using an Agilent 2100 Bioanalyzer. Following quality assessment, sequencing libraries were constructed by fragmenting cDNA, performing end-pair and A-tailing and ligating adapters compatible with the sequencing platform.

Sequencing was performed on an Illumina NovaSeq platform using the S4 flow cell platform in a paired-end format. Read 1 (28 bp) was used to sequence the transcript barcode, while Read 2 (91 bp) captured the transcript sequence. Unique Index read pairs (i7: 8 bp, i5: 8 bp) were used for demultiplexing. The sequencing was covered for an average depth of 40,000 reads per cell. Raw sequencing data were processed using Fluent Biosciences’ PIPseq analysis pipeline (v3.2). Reads were aligned to the GRCm39 reference genome, and unique molecular identifiers (UMIs) were used to deduplicate transcripts. Cells with fewer than 500 detected genes, greater than 20% mitochondrial transcript counts, or fewer than 1000 UMI counts were excluded from further analysis. Quality control and visualization were performed using Seurat (v4.3.0). Data were SCTransformed and scaled using Seurat. Highly variable genes were identified, and dimensionality reduction was performed using principal component analysis (PCA). The first 20 principal components were used to compute a uniform manifold approximation and projection (UMAP) for visualizing the dataset. Clustering was performed using the Louvain algorithm implemented in Seurat to identify groups of cells with similar expression profiles based on shared nearest neighbors in the principal component space. It was further optimized iteratively to capture biologically meaningful subpopulations across different scales. After initial clustering, sub-clustering was performed on individual groups to further resolve heterogeneous populations. The clusters and subclusters were visualized using UMAP to facilitate interpretation and identification of distinct cell populations. Differential expression analysis was performed to compare gene expression levels between clusters or predefined groups of cells. The FindMarkers function in Seurat was used with the Wilcoxon rank-sum test, a non-parametric method that evaluates differences in gene expression without assuming a normal distribution. For statistical significance, genes were considered differentially expressed if they had an adjusted p-value < 0.05 (calculated using the Benjamini-Hochberg method) and a log2 fold change > 1. Data from multiple datasets were merged into a single data frame in R using the merge function, combining shared gene expression features. After integration, the merged dataset was normalized and scaled using Seurat to account for differences in sequencing depth and technical variability. Batch effects were corrected using Harmony (v1.0), which ensured effective alignment of datasets while preserving biologically meaningful variation.

### Flow cytometry and antibodies

Flow cytometry was performed using Cytek Aurora spectral flow cytometer. Single cell suspensions were prepared from thymus, spleen and tumor infiltrating lymphocytes. Cells were first resuspended with Ghost Dye V510 (Cat # SKU 13-0870-T100, Cytek Biosciences) in PBS at room temperature for 20 minutes. After that, cells were stained with cell surface mouse antibodies based on the needs of experiments including PerCP/Cyanine5.5 anti-mouse TCRβ chain Antibody (Biolegend, Cat#109227, Clone H57-597), BUV395 Rat Anti-Mouse CD4 (BD Biosciences, Cat#563790, clone GK1.5), BUV496 Rat Anti-Mouse CD8a (BD Biosciences, Cat#569181, Clone 53-6.7), Brilliant Ultra Violet™ 563 CD25 monoclonal antibody (Thermofisher, Cat# 365-0251-82, Clone PC61.5), BUV615 Rat Anti-Mouse CD24 (BD biosciences, Cat#751499, Clone M1/69), BV711 Rat Anti-Mouse CD11a (BD biosciences, Cat#740676, Clone M1/4), BV570 anti-mouse CD44 Antibody (Biolegend, Cat#103037, Clone IM7), PE/Cy7 anti-mouse CD62L Antibody (Biolegend, Cat#104418, Clone, MEL-14), APC/Cy7 anti-mouse PD-1 Antibody (Biolegend, Cat#135224, Clone 29F.1A12), BV785 anti-mouse antibody (Biolegend, Cat#104543, Clone H1.2F3), BV750 Rat Anti-Mouse CD117 (BD Biosciences, Cat#747412, Clone 2B8), BV650 anti-mouse CX3CR1 (Biolegend, Cat#149033, Clone SA011F11), BUV737 Rat Anti-Mouse CD127 (BD Biosciences, Cat#612841, Clone SB/199), PE/Fire 810 anti-mouse Tim-3 (Biolegend, Cat#149033, Clone RMT3-23) resuspended in the FACS buffer (PBS + 2% FBS + 2mM EDTA). Other surface mouse antibodies for gating out lineage negative populations in thymus including BUV805 anti-mouse CD11b (eBioscience, Cat#368-0112-82, Clone M1/70), BV605 anti-mouse Gr-1 (Biolegend, Cat#108440, Clone RB6-8C5), PE/Fire 700 anti-mouse NK1.1(Biolegend, Cat#156528, Clone S17016D), cFluor UV440 anti-mouse CD19 (Cytek Biosciences, Cat# SKU R7-20835, Clone 6B5), RB613 Rat Anti-Mouse Ly-76 (BD Biosciences, Cat #759470, Clone TER-119), RB780 Hamster Anti-Mouse CD11c (BD Biosciences, Cat #755338, Clone HL3) and FITC anti-mouse TCR γ/δ Antibody (Biolegend, Cat#107504, Clone UC7-13D5). For intracellular staining, cells were permeabilized using Fixation/Permeabilization Concentrate (Cat#00-5123-43, eBioscience) and Fixation/Perm Diluent (Cat#00-5223-56, eBioscience) in the ratio of 1:3 for 20 minutes or overnight at 4°C. After that, cells were then washed using diluted 1X FoxP3 washing (Perm) buffer (Permeabilization Buffer 10X, Cat# 00-8333-56, Invitrogen) and stained with intracellular antibodies. The rabbit polyclonal antibody to mouse NKG7 (UniProtKB – Q99PA5) was obtained by immunizing a rabbit with keyhole limpet hemocyanin-conjugated NKG7 peptide DFWIVATGPHFSAHSGLWPTSQET (Cocalico Biologicals, Reamstown, PA). The anti-NKG7 polyclonal rabbit serum was affinity purified using Sulfolink (Cat#20401, Thermo Fisher Scientific, Waltham, MA) as per manufacturer’s instructions. NKG7 antibody was then labeled with Alexa Fluor™ 594 antibody labeling (Cat# A20185, Invitrogen) based on manufacturer instructions. Other intracellular mouse antibodies including RB545 anti-Tbet (BD biosciences, Cat#569253, Clone O4-46), BUV395 anti-Eomes (BD biosciences, Cat#567171, Clone X4-83), PE anti-Granzyme B Recombinant Antibody (Biolegend, Cat#396406, Clone QA18A28), BV421 anti-Perforin antibody (Biolegend, Cat#154319, Clone S16009A), BUV661 Hamster anti-KLRG1(BD biosciences, Cat#741586, Clone 2F1), anti-TOX Antibody-REAfinity (Miltenyl Biotech, Cat#130-118-335, Clone REA473), anti-TCF1 antibody (Cell signaling technology, Cat#9066S, Clone C63D9). For CD107a degranulation and IFN-γ assay, isolated thymocytes or spleen cells were co-cultured with diluted 1X brefeldin (Biolegend, Cat#420601) and 1X monensin (Biolegend, Cat#420701) with APC anti-mouse CD107a antibody (Biolegend, Cat#505809, Clone XMG1.2) and stimulated with PMA (50ng/mL, Sigma, Cat#P1585) and ionomycin (500 ng/mL, Sigma, Cat#I0634) for 5 hours at 37°C before live and dead cell staining. Data acquisition was conducted using the Aurora’s full-spectrum detection system. Compensation and spectral unmixing were performed using SpectroFlo software (Cytek Biosciences). Gating strategies were applied using FlowJo software (v10.0.0). Fluorescence-minus-one (FMO) controls and unstained controls were included for accurate gating and to validate antibody specificity.

### Spatial transcriptomics of Human Bladder cancer tissue

Human muscle-invasive bladder cancer tissue slides were selected with the approval of Mayo Clinic IRB# 21-000078. These patients underwent radical cystectomy, but did not receive neoadjuvant therapies. Tissue sectioning and immunohistochemical (IHC) staining were conducted at the Pathology Research Core (Mayo Clinic, Rochester, MN) using the Leica Bond RX automated stainer (Leica). Formalin-fixed, paraffin-embedded (FFPE) tissue sections (5 μm thick) were mounted on glass slides and processed for GeoMx DSP. Tissue sections were deparaffinized, rehydrated, and subjected to antigen retrieval before incubation with fluorescently labeled morphology markers to visualize regions of interest (ROIs). Slides were incubated with rabbit anti-human NKG7 monoclonal antibody (1:1000, Dong lab) and anti-human CD8 antibody (1:500, RIV11 clone, Thermo Scientific, MA5-48276), followed by Goat anti-Rabbit IgG (H+L) Cross-Adsorbed Secondary Antibody, Alexa Fluor 532 (Thermofisher Scientific, Cat#A-11009) and Goat anti-Mouse IgG (H+L) Cross-Adsorbed Secondary Antibody, Alexa Fluor 647 (Thermofisher Scientific, Cat#A-11009). ROIs were selected based on fluorescence imaging using the GeoMx instrument. The selected ROIs were then hybridized with barcoded RNA detection probes (GeoMx DSP RNA assays). Following hybridization, ultraviolet (UV) light was used to release ROI-specific oligonucleotide barcodes. Released barcodes were collected, amplified by PCR. Library quality and quantity were assessed using an Agilent 2100 Bioanalyzer and qPCR. Sequencing was performed on an Illumina NovaSeq S4 platform, generating paired end reads. Data analysis was conducted using the GeoMx NGS Pipeline software for spatial alignment, normalization, and quantification of transcript counts. Further analyses, including ROI-specific differential gene expression and pathway enrichment, were performed using the GeoMx DSP Analysis Suite.

## Acknowledgements

We thank all members of the Dong laboratory for helpful discussions and critical analysis of the manuscript. We appreciate Cindy Liu and Susan Harrington for animal breeding and care. We appreciate Roxane Lavoie for assisting in immunofluorescent staining and Yohan Kim for assisting labeling NKG7 antibody. We acknowledge the Genome Analysis Core at Mayo Clinic, including technologists Vernadette A. Simon and Fariborz Rakhshan-Rohakhtar as well as supervisors Julie S. Lau, Samantha J. McDonough, and Mark Mutawe. We are grateful for the spatial transcriptome service provided by Zhenglong Liu, Will Sherman, Christy Trussoni as well as center director Dr. Tamas Ordog. BioRender was used to create the graphics throughout the manuscript. This study was partially supported by NIH grant R01 CA256927 (HD), the Eric & Wendy Schmidt Fund (HD) and Scholarship of Mayo Clinic Graduate School of Biomedical Sciences (ZM).

## Author contributions

Conceptualization: Z.M. and H.D.; Methodology: Z.M. and H.D.; Investigation: Z.M., J.B.H., J.K.G., M.M., M.A.H., E.R.D., W.Z., J.J.T., Y.L., G.Z., F.L., H.BdS., D.D.B. and H.D.; Visualization: Z.M. and H.D.; Funding acquisition: H.D.; Project administration: H.D. and Z.M.; Supervision: H.D.; Writing-original draft: Z.M. and H.D. Writing-review & editing: Z.M., J.B.H., J.K.G., M.M., M.A.H., E.R.D., W.Z., J.J.T., Y.L., G.Z., F.L., H.BdS., D.D.B. and H.D.

## Conflict of interests

H.BdS. is an advisor for the International Genomics Consortium. The remaining authors declare no competing interests.

## Data and materials availability

All materials and data are available from the corresponding author upon request. All data needed to evaluate the conclusions in the paper are present in the paper and/or the Supplementary Materials.

---

**Supplementary Fig 1.**
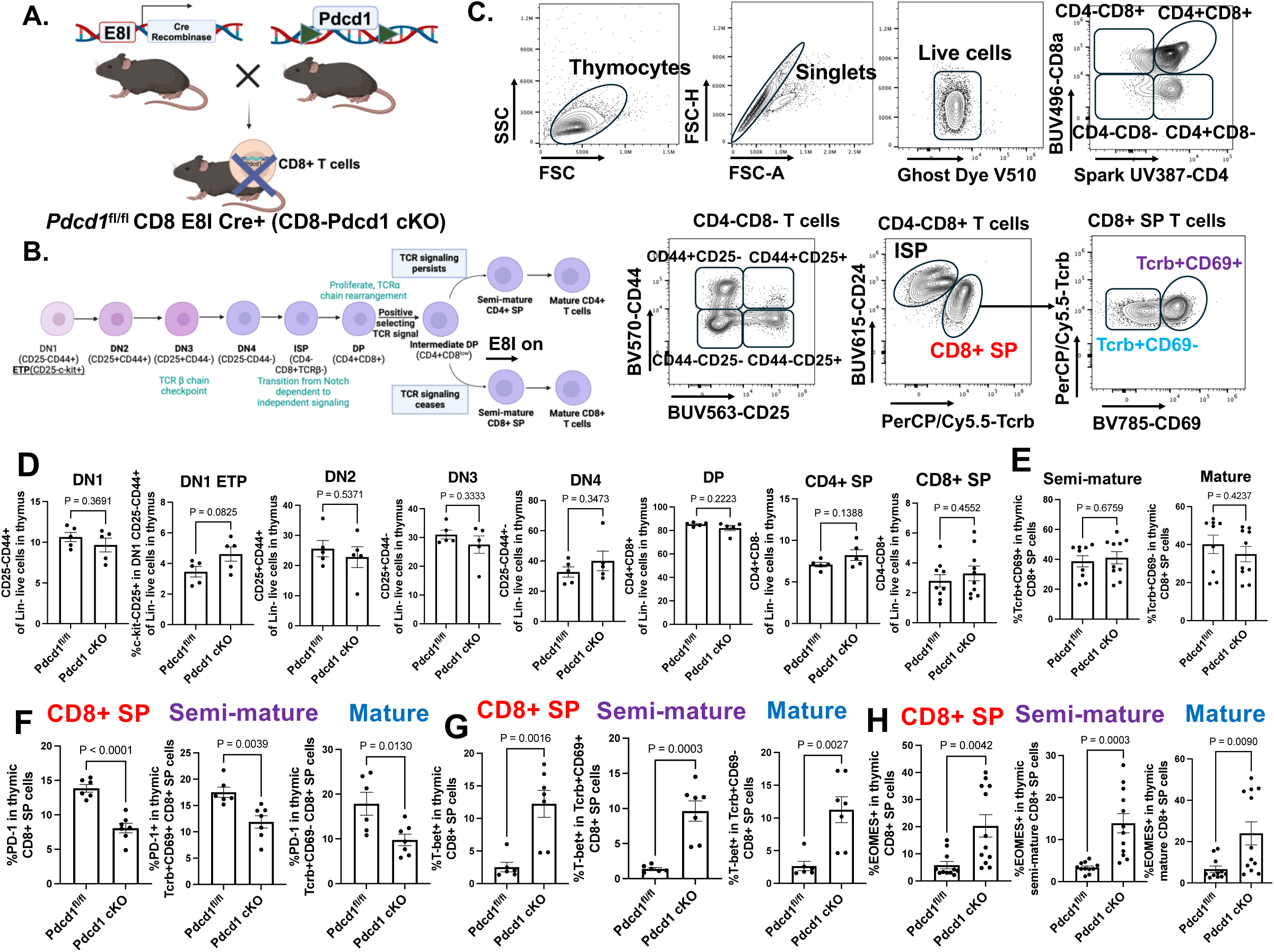
Flow cytometry analysis of the thymic profile of single positive CD8+ T cells. (**A**) Schematic of the production of *Pdcd1*^fl/fl^ CD8 E8I Cre+ (CD8-Pdcd1 cKO) mice. **(B)** Schematic of the stage in which E8i enhancer works during thymic T cell development. (**C**) Flow cytometry analysis of thymic post-positively selected single positive (SP), semi-mature (TCRb+CD69+) and mature (TCRb+CD69-) SP CD8+ T cells. ISP: immature single positive. (**D**) Frequency of double negative (DN)1, DN1 ETP (early T lineage precursor), DN2, DN3, DN4, double positive (DP), CD4+ SP and CD8+ SP thymocytes. (**E**) Frequency of thymic semi-mature and mature SP CD8+ T cells in all SP CD8+ thymocytes. (**F**-**H**) Frequency of PD-1+ (**F**), T-Bet+ (**G**) and EOMES+ (**H**) in thymic SP CD8+ cells, semi-mature and mature CD8+ SP cells. P value was calculated using an unpaired, two-tailed t test.

**Supplementary Fig 2.**
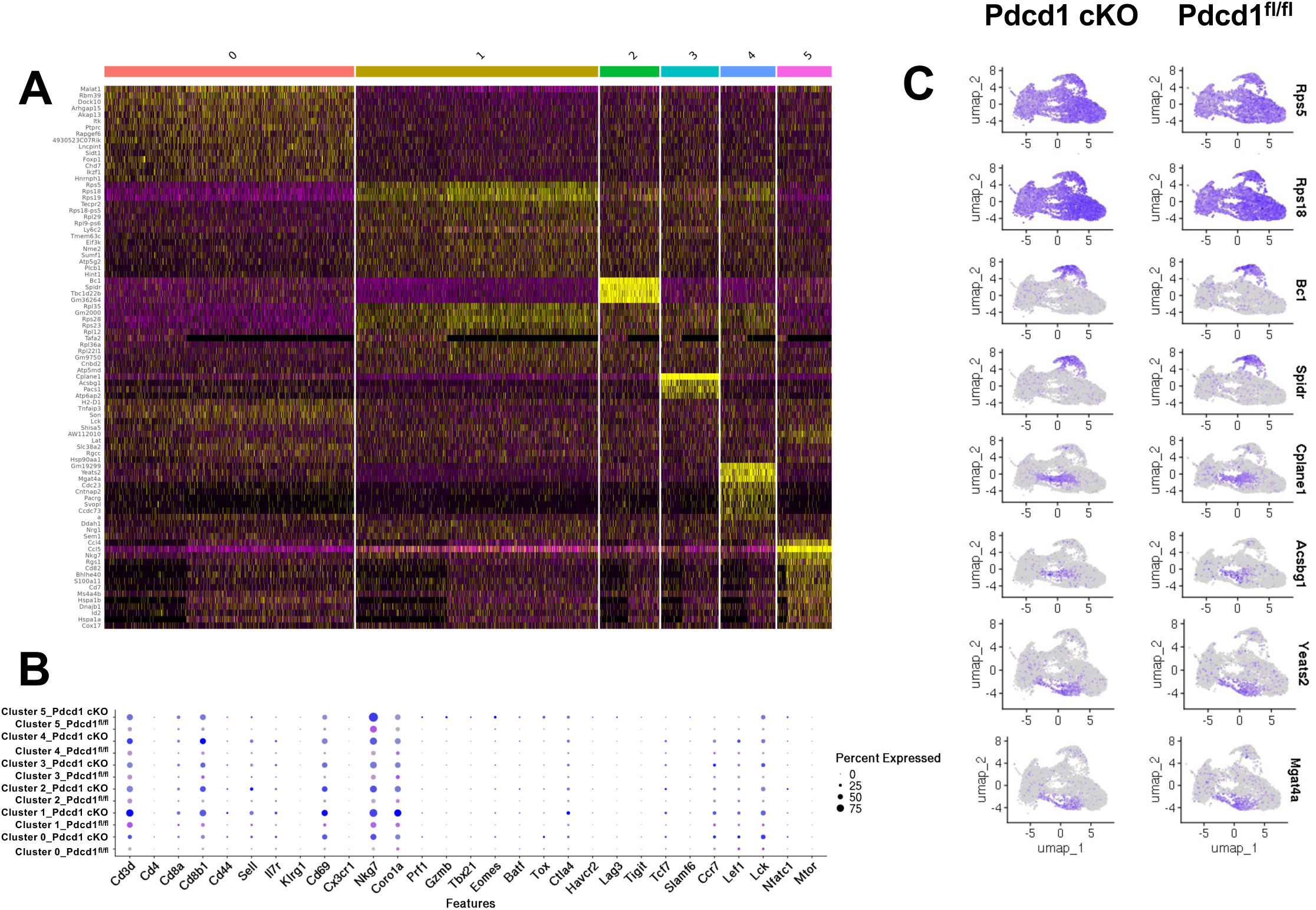
Single cell RNA-seq analysis of splenic CD8+ T cells. (**A**) Heatmap of the differential gene expression of each cluster as shown in the UMAP of splenic CD8+ T cells in Fig. 5A. (**B**) Dotplot graph shows the expression of other genes encoding cytotoxic and exhaustion-associated molecules in splenic CD8+ T cells stratified by control and CD8-Pdcd1 cKO mice. (**C**) Feature plots show the UMAP distribution and expression levels of other featured gene within each cluster stratified by CD8+ T cells isolated from control and CD8-Pdcd1 cKO mice.

**Supplementary Fig 3.**
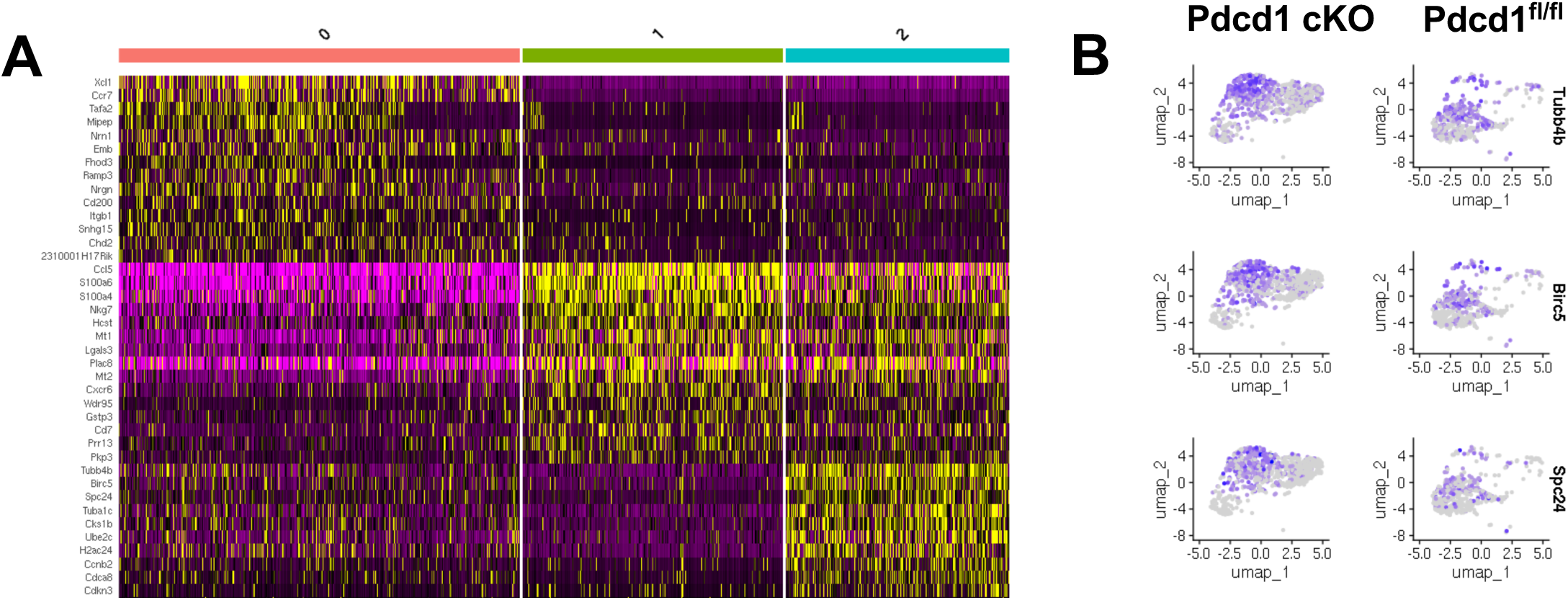
Single cell RNA-seq analysis of CD8+ TILs. (**A**) Heatmap of the differential gene expression of each sub-cluster identified in the UMAP of CD8+ TILs in Fig. 6M**. (B)** Feature plots show the UMAP distribution and expression levels of other featured genes that were only upregulated in cluster 1 stratified by CD8+ T cells from control and CD8-Pdcd1 cKO mice.

**Supplementary Fig 4.**
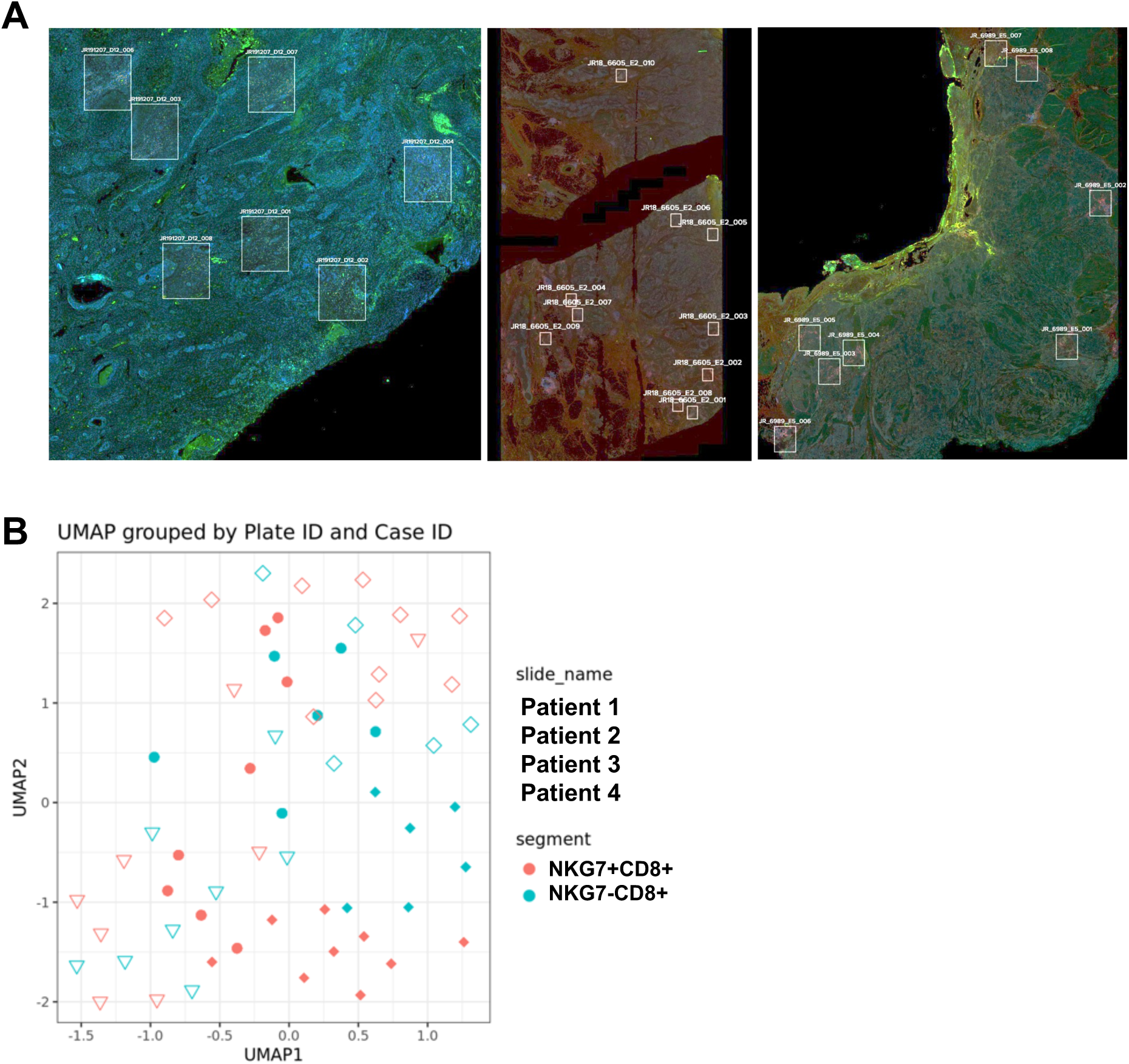
Spatial transcriptome analysis of NKG7 expressing CD8 T cells in human tumor tissues. **(A)** Region of interest selected in tumor slides of other three patients with invasive bladder cancer for spatial transcriptomics in addition to the representative Fig. 7A**. (B)** UMAP graph shows the principal component analysis (PCA) of the four patient tissue samples, highlighting distinct clustering of NKG7+CD8+ vs. NKG7-CD8+ T cells.

**Supplementary Fig 5.**
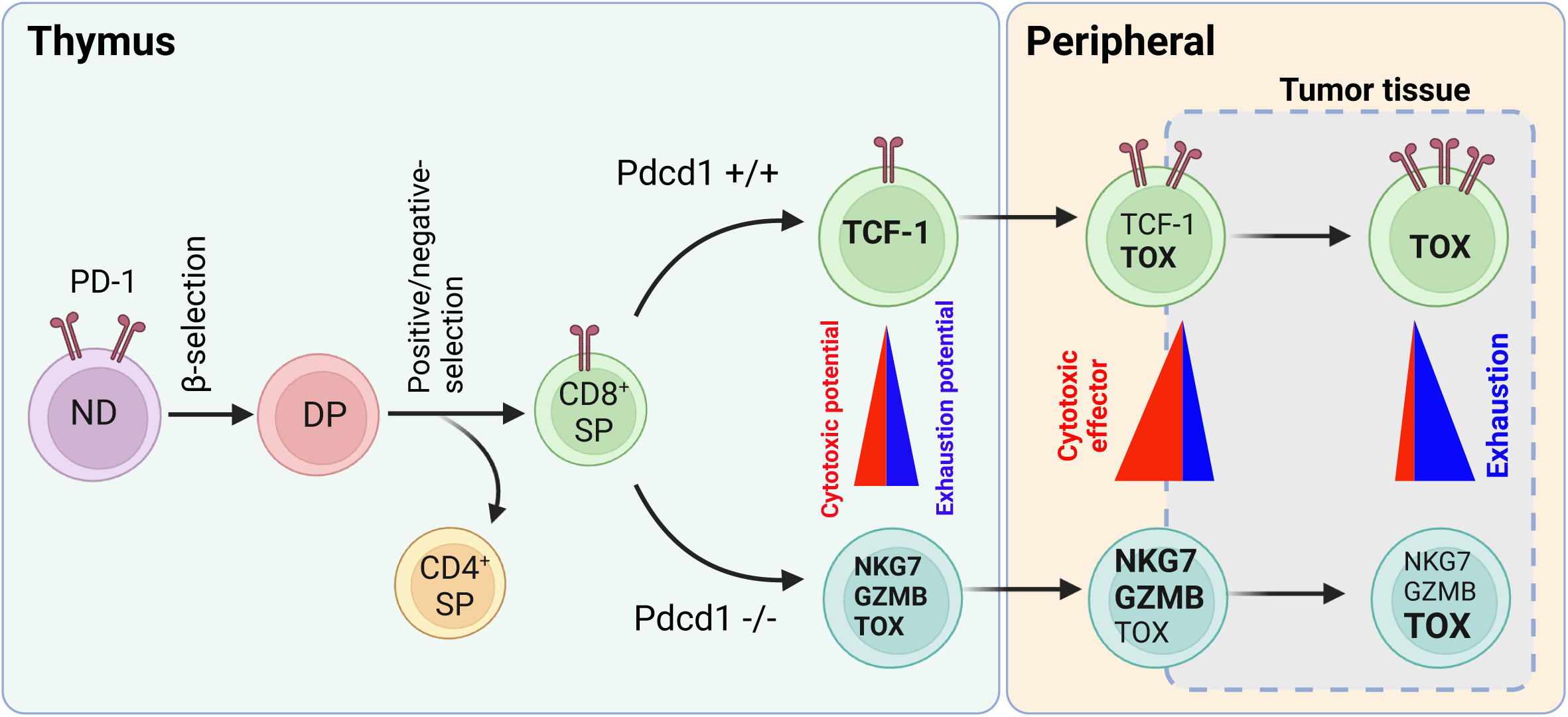
A graphic summary. Schematic role of PD-1 in balance of cytotoxic and exhaustion potential and capacity of CD8+ T cells in thymus and peripheral tissues including tumor tissues.

